# Evolution of gene expression after whole-genome duplication: new insights from the spotted gar genome

**DOI:** 10.1101/151944

**Authors:** Jeremy Pasquier, Ingo Braasch, Peter Batzel, Cedric Cabau, Jérome Montfort, Thaovi Nguyen, Elodie Jouanno, Camille Berthelot, Christophe Klopp, Laurent Journot, John H. Postlethwait, Yann Guiguen, Julien Bobe

## Abstract

Whole genome duplications (WGD) are important evolutionary events. Our understanding of underlying mechanisms, including the evolution of duplicated genes after WGD, however remains incomplete. Teleost fish experienced a common WGD (teleost-specific genome duplication, or TGD) followed by a dramatic adaptive radiation leading to more than half of all vertebrate species. The analysis of gene expression patterns following TGD at the genome level has been limited by the lack of suitable genomic resources. The recent concomitant release of the genome sequence of spotted gar (a representative of holosteans, the closest lineage of teleosts that lacks the TGD) and the tissue-specific gene expression repertoires of over 20 holostean and teleostean fish species, including spotted gar, zebrafish and medaka (the PhyloFish project), offered a unique opportunity to study the evolution of gene expression following TGD in teleosts. We show that most TGD duplicates gained their current status (loss of one duplicate gene or retention of both duplicates) relatively rapidly after TGD (i.e. prior to the divergence of medaka and zebrafish lineages). The loss of one duplicate is the most common fate after TGD with a probability of approximately 80%. In addition, the fate of duplicate genes after TGD, including subfunctionalization, neofunctionalization, or retention of two ‘similar’ copies occurred not only before, but also after the radiation of species tested, in consistency with a role of the TGD in speciation and/or evolution of gene function. Finally, we report novel cases of TGD ohnolog subfunctionalization and neofunctionalization that further illustrate the importance of these processes.

## Introduction

Whole genome duplication (WGD) events have played important roles in the evolution of many living organisms, including vertebrates. Two WGD events (VGD1 and VGD2) occurred at the root of the vertebrate radiation as initially postulated by Ohno (Ohno, 1970; Dehal and Boore, 2005; Nakatani et al., 2007; Canestro et al., 2009). Another WGD event (the Teleost-specific genome duplication or TGD) occurred at the root of the teleost lineage (Amores et al., 1998; Postlethwait et al., 1999; Taylor et al., 2003; Jaillon et al., 2004) and was followed by an important adaptive radiation (Glasauer and Neuhauss, 2014). With more than 30,000 species, living teleost fish occupy a wide diversity of aquatic habitats, including the most extreme ones, like deep-sea vents, frigid Antarctic waters, acid hot springs, and ephemeral pools. The relation of the TGD and teleost biodiversity, however, remains intricate and is uncoupled in geological time (Santini et al., 2009; Clarke et al., 2016). After genome duplication by autopolyploidy, both duplicated gene copies initially encode proteins with identical sequences and expression patterns; over time, some gene pairs revert to single copy (loss of one of the duplicate copies), while members of other pairs evolve new functions (neofunctionalization), including different tissue-specific expression domains, or share between the two duplicates the functions of the ancestral single copy gene (subfunctionalization), or both (Force et al., 1999; He and Zhang, 2005). Evidence also indicates that both copies are sometimes retained to produce enough of the proteins to perform the same ancestral function (retention for dosage constraint called quantitative subfunctionalization (Force et al., 1999)). In teleost fish, the frequency of gene loss following the TGD was recently studied using genomes from phylogenetically distant lineages (Inoue et al., 2015). The process has however not yet been studied using a close outgroup for the TGD. The gar (*Lepisosteus oculatus*) genome provides, for the first time, a representative of the most recently diverging lineage before the TGD to help evaluate evolution of gene expression after the TGD.

## Material and Methods

### Gene dataset: linking spotted gar genes to TGD ohnologs and singletons in zebrafish and medaka

#### Identification of TGD ohnologs in zebrafish and medaka and their single gar ortholog

To identify orthologs of spotted gar protein-coding genes, zebrafish (*Danio rerio*) and medaka (*Oryzias latipes*) predicted intragenomic paralogs were downloaded from Ensembl74 along with their spotted gar orthologs using Biomart (Kinsella et al., 2011). For the initial lists of intragenomic paralogs of zebrafish/medaka, we used the Biomart ‘Homologs: Paralogs’ function to obtain the Ensembl gene IDs and genomic locations of Ensembl-predicted paralogs and the ‘Homology Type’ and ‘Ancestor’ information of the predicted duplication; we further used the ‘Homologs: Orthologs’ function to obtain the predicted gar ortholog of each pair of paralogs. These paralogous pairs were filtered for the duplication ancestor Clupeocephala (supercohort Clupeocephala), the most basal duplication point available with whole genome sequences in Ensembl after divergence of teleosts from gar. Next, each paralog in zebrafish/medaka was checked for being present only once in the gene dataset of intragenomic paralogs, thereby leading removing gene duplicates that appeared after the TGD within the lineages leading to zebrafish and medaka, respectively. Furthermore, each gene pair was required to have a unique, single gar ortholog to remove paralog pairs for which no gar ortholog was available or for which gene duplication(s) occurred within the gar lineage. Cases of ‘split genes’ (i.e. genes present twice in the dataset according to Ensembl) were removed as well. This process yielded a total of 1,901 cases of 1:2 gene relations between gar and zebrafish and 1,597 cases of 1:2 gene relationships between gar and medaka. To further filter for zebrafish and medaka paralogs that show the expected pattern of double conserved synteny generated by the TGD, the 1:2 spotted gar vs. zebrafish/medaka gene trios were required to be located in paralogous clusters defined by the Synteny Database (http://teleost.cs.uoregon.edu/syntenydb/) (Catchen et al., 2009) using zebrafish/medaka as source genomes and spotted gar as outgroup genome (sliding window size: 200 genes; membership ≥10 paralogous pairs). After this conserved synteny filtering, 1,606 pairs of zebrafish (Fig. 1A, orange circle) and 1,315 of medaka (Fig. 1A, purple circle) paralogs were retained that are considered a highly stringent curated subset of ‘TGD ohnologs’ (paralogs derived from a genome duplication event, see Fig. 1B) having both phylogenetic and synteny support for their origin in the TGD. Zebrafish and medaka TGD paralog pairs were joined based on their single gar ortholog and orthology of zebrafish genes to medaka genes was confirmed by patterns of medaka/zebrafish conserved synteny obtained with the Synteny Database. A total of 774 TGD paralog pairs shared between zebrafish and medaka were defined (Fig. 1A, orange/purple intersection). For further analysis, the TGD ohnolog list of zebrafish (1,606 pairs) was randomized with respect to the assignment of one or the other TGD ohnologs of a pair as “Ohnolog1” or “Ohnolog2”. Assignment to “Ohnologl” or “Ohnolog2” for the 774 TGD ohnologs shared between zebrafish and medaka followed the randomized zebrafish assignment. The remaining 541 TGD ohnologs from medaka not shared with zebrafish were further randomized as “Ohnolog1” or “Ohnolog2”.

**Figure 1:**
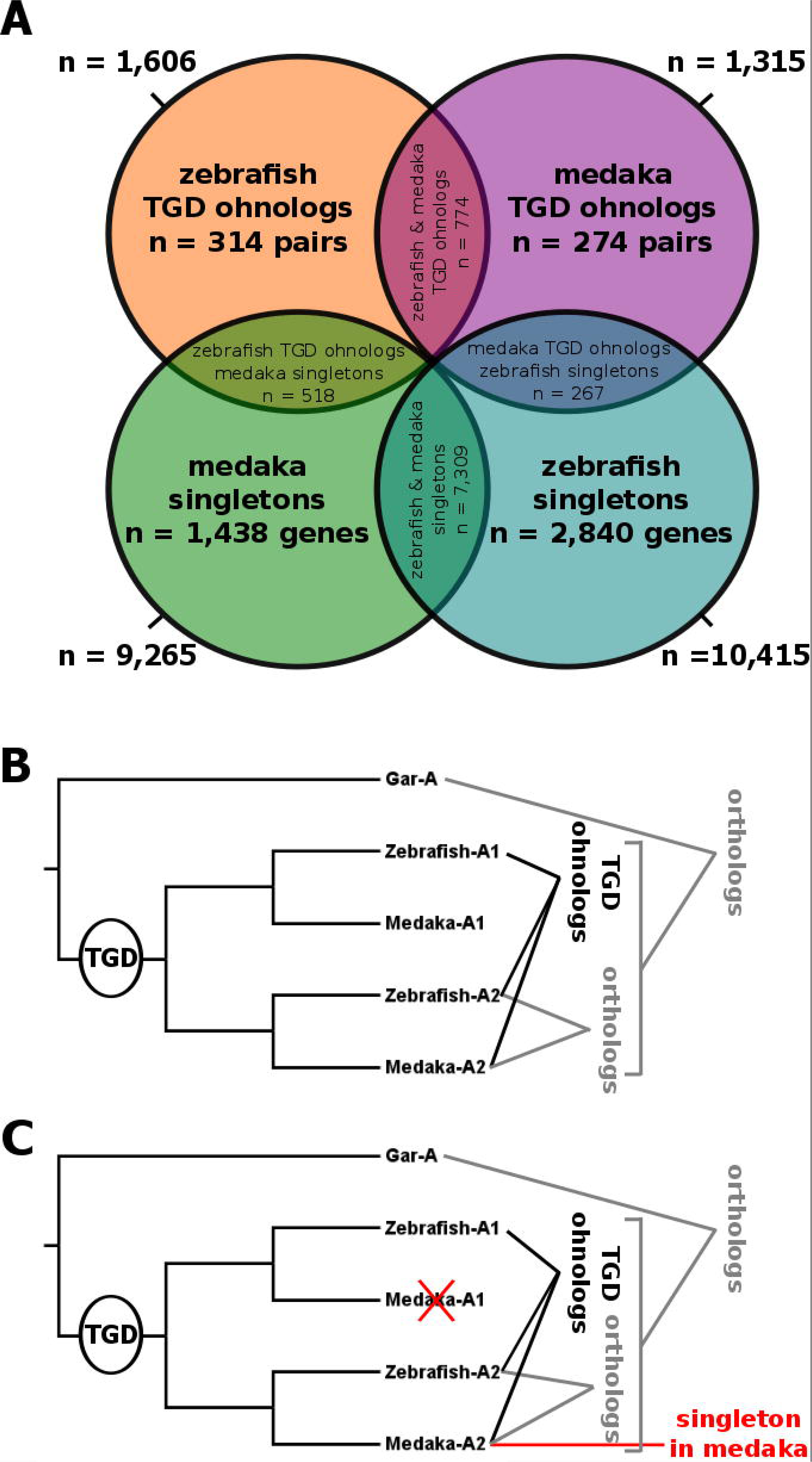
The data set. The diagram (A) shows the partitioning of Gar genes and their teleost orthologs used in the study. Numbers indicate the number of genes based on their presence as singleton or TGD ohnologs in zebrafish and medaka. Orthology and paralogy (including ohnologs, paralogs originating from TGD) relationships are illustrated in panels B and C for ohnolog pairs and singletons, respectively.

#### Identification of zebrafish and medaka singletons

To identify singletons (i.e. genes for which one of the two TGD ohnologs was lost following TGD in zebrafish and/or medaka, see Fig. 1C), we removed genes from the BioMart-derived list of intragenomic paralogs that had an indication for TGD duplication (duplication ancestor Clupeocephala, see above) as well as for lineage-specific gene duplication after the TGD (e.g. for zebrafish, duplication ancestors Otophysi (section Otophysa) and *Danio* (genus *Danio*). We also removed genes with duplication ancestor Neopterygii (subclass Neopterygii, i.e. the ancestor of gar and teleosts) because these inferred duplication nodes could be artifacts of tree reconstructions in Ensembl and thus potentially include TGD ohnologs or other types of gene duplication events that occurred within teleosts. Genes with Ensembl gene names indicative of gene duplication [e.g., zebrafish gene ‘TRIM8 (1 of 4), ENSDARG00000017173; medaka gene ‘KIAA1598 (3 of 3)’, ENSORLG00000011511] were removed from the list of singletons as well. Each zebrafish or medaka singleton gene was required to have a unique, single gar ortholog (‘ortholog-one-to-one’) to remove genes for which no gar ortholog was available or for which gene duplication(s) occurred within the gar lineage. Genes located on unplaced scaffolds or mitochondrial genomes in zebrafish/medaka were removed as well. This survey left us with a list of 10,415 and 9,265 genes in zebrafish (Fig. 1A, green circle) and medaka (Fig. 1A, turquois circle), respectively, with a 1:1 relationship to a single gar gene, thus, genes likely to be singletons with respect to the TGD.

To detect shared singleton genes in zebrafish and medaka, the lists of singletons of both species were joined based on their single gar ortholog, identifying a subset of 7,309 genes being singletons in both zebrafish and medaka (Fig. 1A, green/turquois intersection), suggesting that the second TGD ohnolog of these genes was lost before the divergence of the zebrafish and medaka lineages, in other words, relatively early during teleost evolution within a few tens of millions of years following the TGD (Broughton et al., 2013).

Finally, remaining (i.e., not shared) singletons of one teleost species (zebrafish/medaka) were joined with TGD ohnolog lists of the other species (medaka/zebrafish) based on their single gar ortholog. This process led to an intersection of 267 zebrafish singletons with medaka TGD ohnolog pairs (Fig. 1A, turquois/purple intersection), and 518 medaka singletons that merged with zebrafish TGD ohnolog pairs (Fig. 1A, green/orange intersection). The singleton gene of one species was assigned orthologous to “Ohnolog1” or “Ohnolog2” of the other species based on patterns of conserved synteny obtained from the Synteny Database (Catchen et al., 2009) (Fig. 1B).

#### Biological material and tissue-specific RNA-seq libraries

RNA-seq data used in the present study for spotted gar, zebrafish, and medaka originated from the public PhyloFish database (http://phylofish.sigenae.org/index.html). Extensive description of the biological samples used, RNA-seq methods, and *de novo* assemblies of gene repertoires can be found in the original report of the PhyloFish database (Pasquier et al., 2016) and the gar genome publication (Braasch et al., 2016). Briefly, tissue-specific transcriptomes were generated in each species using the following tissues: ovary, testis, brain, gills, heart, muscle, liver, kidney, bone, intestine, and embryos at the stage that eyes first become pigmented. For each tissue of a given species, a single library was constructed. Multiplexed paired-end (2 × 100 bp) sequencing was performed using an Illumina HiSeq2000 instrument with a minimum of 40 million reads per library. For all species, tissues were sampled from the same female individual and testis from a male individual, when possible. In some species and depending on the tissues, RNA samples from different individuals were pooled to obtain sufficient amounts of RNA for sequencing. All corresponding information is available in the biosample and bioproject files deposited in SRA under BioProject accession # PRJNA255889 (medaka), PRJNA255848 (zebrafish) and PRJNA255881 (gar). Corresponding data were deposited into NCBI SRA database under accession numbers: SRP044781 (zebrafish), SRP044784 (medaka), and SRP044782 (gar). For medaka and zebrafish, all sampled fish originated from the INRA LPGP experimental facility. Fish were reared and handled in strict accordance with French and European policies and guidelines of the INRA LPGP Institutional Animal Care and Use Committee (# 25M10), which approved this study. In medaka, all tissues were collected from 11-month old fish, with the exception of ovary and testis that were collected from 2-month old fish. In zebrafish, all samples were collected from 2-month old fish. For gar, adult tissues were collected from wild animals in Louisiana. Embryos were grown at the University of Oregon.

#### Gene expression patterns using RNA-seq reads

To study expression patterns and levels of zebrafish, medaka, and spotted gar transcripts, a reference coding sequence (CDS) library was built for each species. Each library was deduced from the zebrafish (assembly Zv9), medaka (assembly MEDAKA1) and gar (assembly LepOcu1) Ensembl genomic databases as follows: for each gene, one CDS was retained in the library; when multiple CDS were referenced for a single gene, the longest CDS was retained as representative of the gene product. We then mapped our double stranded RNA-seq reads onto the corresponding CDS library using BWA-Bowtie (Li and Durbin, 2009) with stringent mapping parameters (maximum number of allowed mismatches –aln 2). Mapped reads were counted using SAMtools (Li et al., 2009) idxstat command, with a minimum alignment quality value (−q 30) to discard ambiguous mapping reads. For each species, the numbers of mapped reads were then normalized for each gene across the 11 tissues using DESeq (Anders and Huber, 2010).

#### Evolution of gene expression after TGD in zebrafish and medaka

To compare expression patterns of genes that have been retained as singletons after the TGD to the expression pattern of TGD ohnologs, we created an average expression pattern for each pair of ohnologs by calculating, individually for each of the 11 tissues, the average expression level between the two ohnologs. This average expression pattern is designated as ‘ohnolog pair’ (or ohno-pair). Using Pearson’s correlation in R, we determined the expression pattern correlation between each zebrafish or medaka gene to its gar ortholog. We then performed a multiple two-sided Wilcoxon Mann-Whitney test to compare the mean correlation of singletons, ohnolog-1, ohnolog-2 and ohnolog-pair within and across species.

To study the relative expression levels of those genes, we calculated the average expression level of each gene over the 11 tissues. We then calculated the ratio of those average expression levels between each zebrafish or medaka gene and its gar ortholog. We performed a multiple two-sided Student t-test to compare the mean expression level ratio of singletons, ohnolog-1, ohnolog-2 and ohnolog-pair within and across species.

To specifically study the evolution of the genes that have been retained as TGD ohnologs in zebrafish and medaka, we determined the expression pattern correlation between ohnolog-1 and ohnolog-2 in each species and between zebrafish ohnologs and their medaka orthologs. We then performed a multiple two sided Wilcoxon Mann-Whitney test to compare the mean correlation of zebrafish ohnologs, medaka ohnologs, zebrafish and medaka orthologs-1, and zebrafish and medaka orthologs-2.

To further study genes that have been retained as TGD ohnologs in zebrafish and medaka, we delineated, for each species, four groups of ohnologs based on (i) correlation between the expression patterns of ohnolog-1 and ohnolog-2 (HC: high correlation, p > 0.05; NC: no correlation, p < 0.05, Pearson’s correlation test), and (ii) the relative expression levels of ohnolog-1 vs. ohnolog-2 (SE: same expression levels, p > 0.05; DE: different expression levels, p < 0.05, Student’s t-test). All tests were performed using R, and a Bonferroni correction was applied on all multiple tests.

#### Clustering analysis

The expression profiles of conserved TGD ohnologs in zebrafish and medaka were also analyzed using supervised clustering (i.e. the order of the samples on the heat maps being similar for all species/analyses). Hierarchical clustering was processed using centroïd linkage clustering with Pearson’s uncentered correlation as the similarity metric on data that were normalized and median-centered using the Cluster program (Eisen et al., 1998). Results (heat maps) of hierarchical clustering analyses were visualized using the Java TreeView program (Eisen et al., 1998).

### Detection of neofunctionalization and subfunctionalization after the TGD in zebrafish and medaka

The calculated Pearson’s correlation between expression patterns of zebrafish or medaka TGD ohnologs and their gar orthologs were also used to detect automatically neofunctionalization and subfunctionalization processes. An r value threshold of 0.75 was used to identify correlated expression profiles. Criteria used to detect neofunctionalization pattern are presented in Supplemental File 1.

## Results

### Phylogenetic and conserved synteny analyses identify gar genes with TGD ohnologs and singletons in zebrafish and medaka

In both teleost species, the number of TGD singletons was much higher than the number of genes retained in duplicates after TGD (i.e. pairs of TGD ohnologs). In zebrafish, 10,415 singletons and 1,606 pairs of TGD ohnologs were identified (Fig. 1, Supplemental file 2). In medaka, 9,265 singletons and 1,315 pairs of ohnologs were identified (Fig. 1, Supplemental file 2). For 774 gar genes, unambiguous pairs of TGD ohnologs could be identified in both zebrafish and medaka. For 7,309 gar genes, unambiguous singletons could be identified in both zebrafish and medaka. For 518 gar genes, orthologous pairs of unambiguous zebrafish TGD ohnolog pairs and unambiguous medaka TGD singletons could be identified. For 267 gar genes, orthologous unambiguous medaka TGD ohnolog pairs and unambiguous zebrafish TGD singletons could be identified in medaka and zebrafish, respectively. For all remaining gar genes, ambiguities remained for either zebrafish or medaka with regard to their TGD ohnolog or singleton status under our stringent phylogenetic and synteny search criteria. For instance, 2,840 gar genes had orthology relationships with zebrafish singletons while it could not established whether or not this gene was present as an unambiguous TGD ohnolog or singleton in medaka (Fig. 1, Supplemental file 2). The numbers of overlapping categories in Figure 1 is thus very likely an underestimate of the actual number.

#### Identification of zebrafish and medaka genes exhibiting neofunctionalization or subfunctionalization following the TGD

Based on the evolution of expression profiles in zebrafish and medaka, in comparison to gar (N=774), a total of 51 (6.6%) and 6 (0.8%) cases could be identified in which TGD ohnolog pairs exhibited clear signatures of neofunctionalization or subfunctionalization following our criteria, respectively (Supplemental File 3).

#### Neofunctionalization

In neofunctionalization, a gene evolves a novel function not present for the preduplication ancestral gene (Ohno, 1971; Force et al., 1999). A total of 17 gar genes were identified with teleost co-orthologs that exhibited neofunctionalization in both teleosts and 34 in only one of the two teleosts (Supplemental file 3). As previously reported (Braasch et al., 2016), *solute carrier family 1 member 3* (*slc1a3*) exhibited a clear pattern of neofunctionalization, showing evolutionary new expression of one of the duplicated copies in the liver in both zebrafish and medaka (Fig. 2A-C). As an example of neofunctionalization that appears to have occurred after the divergence of the zebrafish vs. medaka lineages, *bicaudal D homolog 1* (*bicd1*) ohnolog-1 in medaka Fig. 2F) exhibited significant expression in heart and ovary that was not found in gar or zebrafish (Fig. 2D, E).

**Figure 2:**
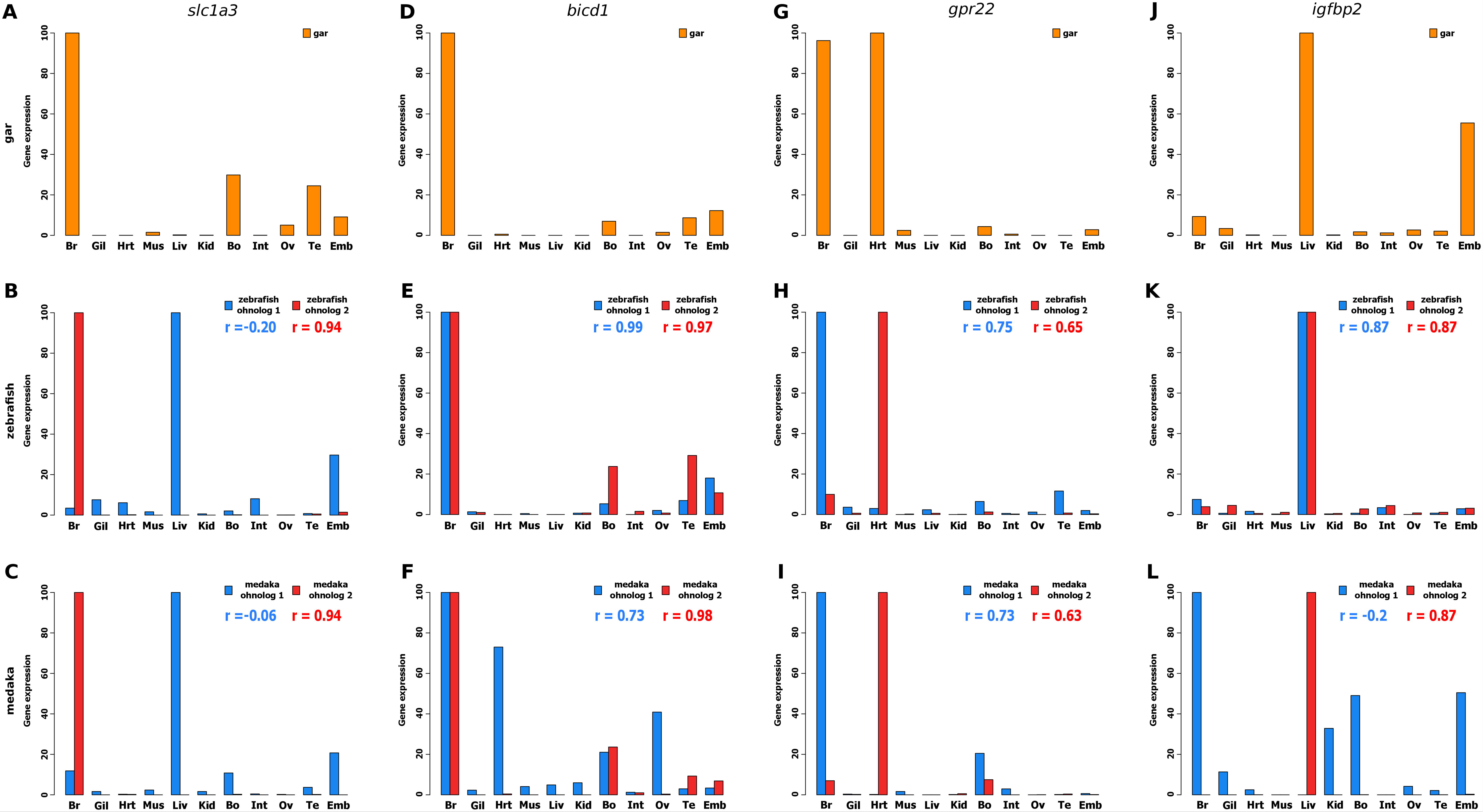
Examples of conserved neofunctionalized and subfunctionalized ohnologs in zebrafish and medaka, based on expression pattern comparisons with their gar orthologs. Examples of neofunctionalized TGD ohnologs: (A-C) *slc1a3:* solute carrier family 1 member 3; (D-F) *bicd1:* bicaudal D homolog 1. Examples of subfunctionalized TGD ohnologs: (G-I) *gpr22*: G protein coupled receptor 22; (J-L) *igfbp2*: insulin-like growth factor binding protein 2. Each graph shows the correlation (r) of gene expression profiles between gar and each of the two ohnologs. For each graph the bars show the expression levels (rpkm, number of reads per kbase per million reads) obtained from the gar, zebrafish, and medaka libraries of the PhyloFish database http://phylofish.sigenae.org/index.html (Pasquier et al., 2016). For all genes, expression is shown in brain (Br), gills (Gil), heart (Hrt), muscle (Mus), liver (Liv), kidney (Kid), bones (Bo), intestine (Int), ovary (Ov), testis (Te), and in a pool of embryos at eyed stage (Emb).

#### Subfunctionalization

In subfunctionalization, functions present in an unduplicated ancestral gene partitioned between the two gene copies after a gene duplication event (Force et al., 1999). A total of five gar genes could be identified with orthologs that exhibited subfunctionalization in both species and one gene showed subfunctionalization in only one species (Supplemental file 3). As previously reported (Braasch et al., 2016), *G protein-coupled receptor 22* (*gpr22*) exhibited clear subfunctionalization of TGD ohnologs with one ohnolog expressed in brain as in gar and the other ohnolog expressed in heart as in gar (Fig. 2G-I). As an example of subfunctionalization occurring after the divergence of zebrafish and medaka lineages, *insulin-like growth factor binding protein 2* (*igfbp2*) was predominantly expressed in liver and embryos in gar, but following the TGD, one medaka ohnolog was predominantly expressed in liver, while the other ohnolog retained strong expression in the embryo, while the zebrafish tended to retain the ancestral pattern in liver for both TGD ohnologs (Fig. 2J-L).

#### Evolution of the expression of singletons and TGD ohnologs following TGD in comparison to gar

In both zebrafish and medaka, the average expression profiles of ohnolog pairs were significantly more correlated to gar than each ohnolog taken separately (Fig. 3A). Similarly, the average expression profiles of ohnolog pairs were significantly more correlated to the expression patterns of their gar ortholog than the expression patterns of teleost singletons were correlated to their gar ortholog (Fig. 3A). No significant difference could be observed between singletons and ohnologs taken separately in either of the two species (Fig. 3A).

**Figure 3:**
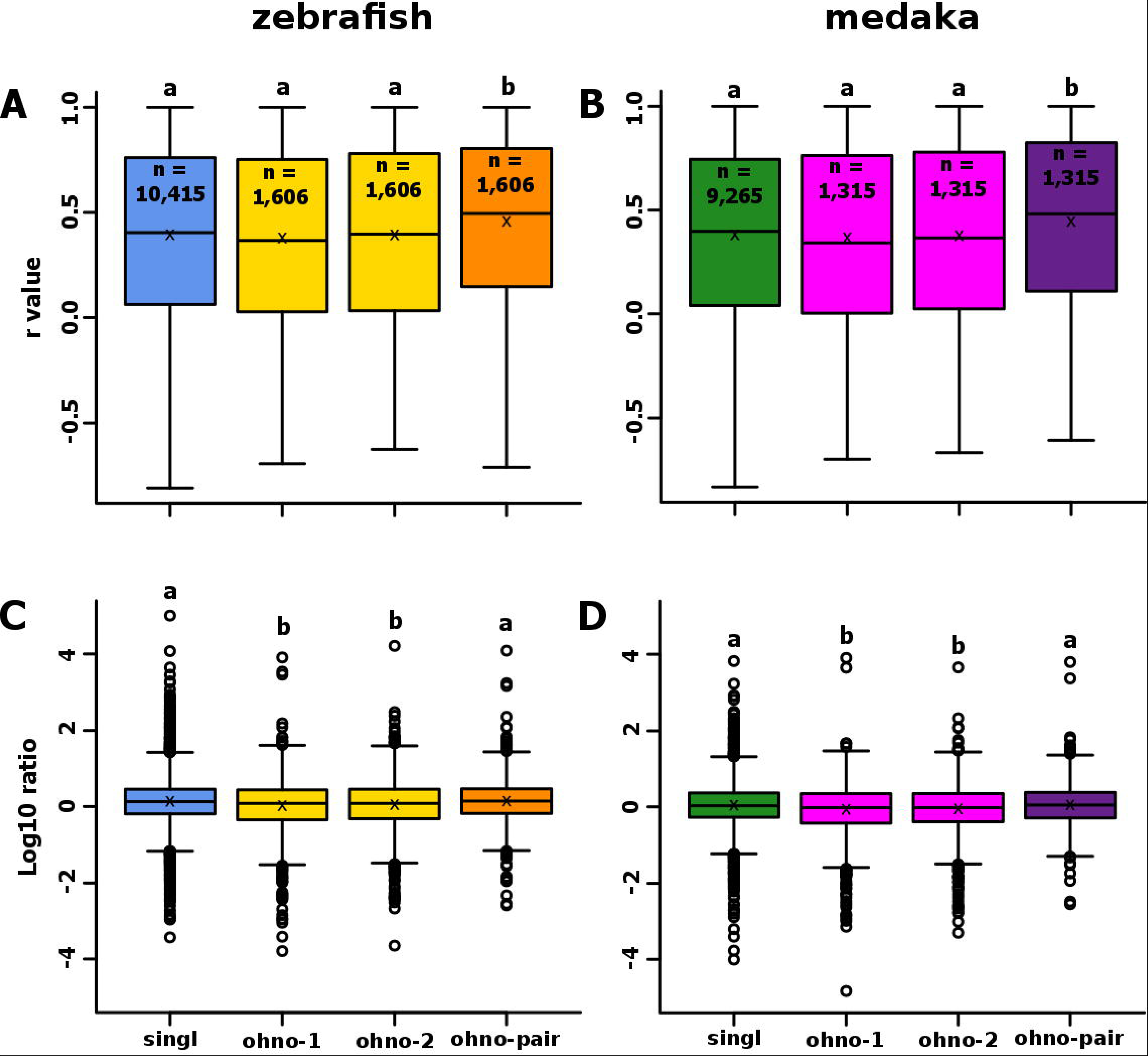
Conservation of expression after the TGD for genes that have been retained in duplicates or as singletons in teleosts. Boxplots representing the distribution of correlations between gene expression patterns of gar genes and their corresponding ortholog(s) in zebrafish (A) or medaka (B). Boxplots representing the distribution of the ratio between the gene expression level of gar genes and their ortholog(s) in zebrafish (C) or medaka (D). For each boxplot, the black line, the black cross, and the black circle represent the median, the mean and the outliers, respectively. ‘Ohno-pair’ represents the average expression profile of a pair of ohnologs as defined in Additional File 1.

In comparison to gar, for the 11 studied tissues, genes retained as singletons following the TGD exhibited a significantly higher average expression level than ohnologous genes taken separately (Fig. 3B), suggesting that expression levels of retained ohnologs tend to decrease so that a duplicated gene pair together approximates the levels of the preduplication gene. In contrast, the average expression level of singletons was not significantly different from the average expression of ohnolog pairs (Fig. 3B). Similar observations were made in both zebrafish and medaka.

#### Evolution of the expression of singletons present in only one species and retained in duplicates in the other species

In the dataset of 518 genes kept as TGD singletons in medaka and as TGD ohnologs in zebrafish, no significant differences in correlation with gar were observed between medaka singletons and their zebrafish ortholog (Fig. 4A). In contrast, a higher correlation with gar was observed in medaka singletons in comparison to the non-orthologous zebrafish TGD ohnolog (Fig. 4A). This difference in correlation was not observed when the group of 267 zebrafish singletons were compared to medaka TGD ohnologs (Fig. 4A).

**Figure 4:**
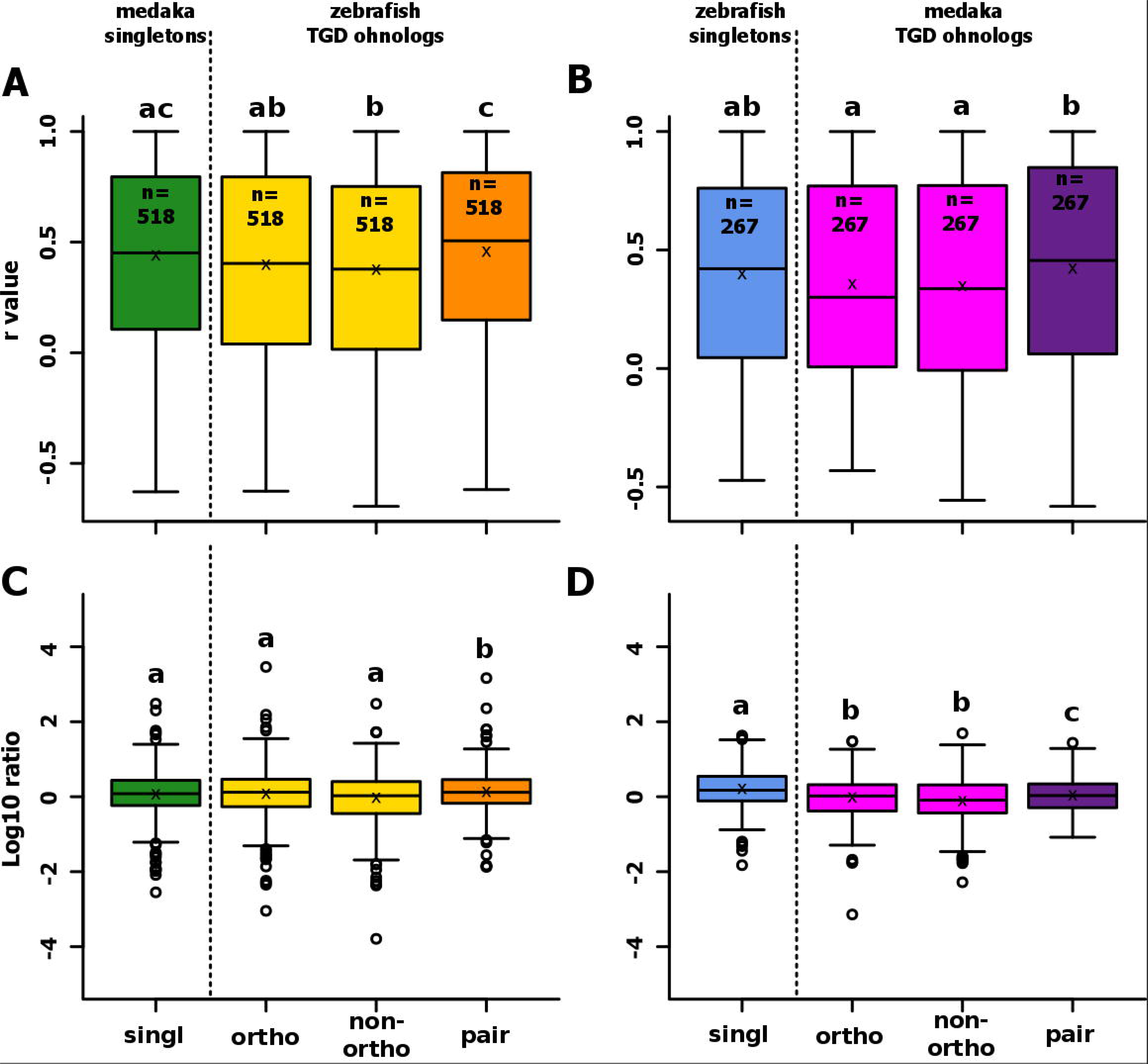
Conservation of expression after the TGD for genes conserved as duplicates in one teleost and as a singleton in the other teleost. Boxplots representing the distribution of correlations between gene expression patterns in gar and their ortholog(s) in zebrafish (A) or medaka (B). Boxplots representing the distribution of the ratio between the gene expression level of genes in gar and their ortholog(s) in zebrafish (C) or medaka (D). For each boxplot, the black line, the black cross, and the black circle represent the median, the mean and the outliers, respectively. ‘Ohno-pair’ represents the average expression profile of a pair of ohnologs as defined in Additional File 1. It needs to be pointed out in the figure that the orthology relations are now different between the two yellow/pink bars to the green/blue one, which is different from Fig. 3.

No difference in expression levels was observed between medaka singleton and both zebrafish TGD ohnologs, orthologous or non-orthologous (Fig. 4B), while a higher expression was observed for zebrafish singletons in comparison to medaka TGD ohnologs. In all comparisons, expression levels in zebrafish vs. medaka shows higher expression in zebrafish. The relatively small number of genes (i.e. N=267) present as singletons in zebrafish and as TGD ohnologs in medaka should be noted, however.

#### Correlation of expression profiles following the TGD for orthologous and ohnologous genes in zebrafish and medaka

Using 774 pairs of TGD ohnologs shared between zebrafish and medaka as defined in Figure 1A (orange/purple intersection), we analyzed the correlation of expression profiles of ohnologs and orthologs. Orthologs exhibited a significantly (p<0.001) higher correlation than ohnologs with correlation (r) of 0.34 and 0.57 for ohnologs and orthologs, respectively (Fig. 5). Similar results were obtained when ohnologs and orthologs of the two species were analyzed separately (data not shown).

**Figure 5:**
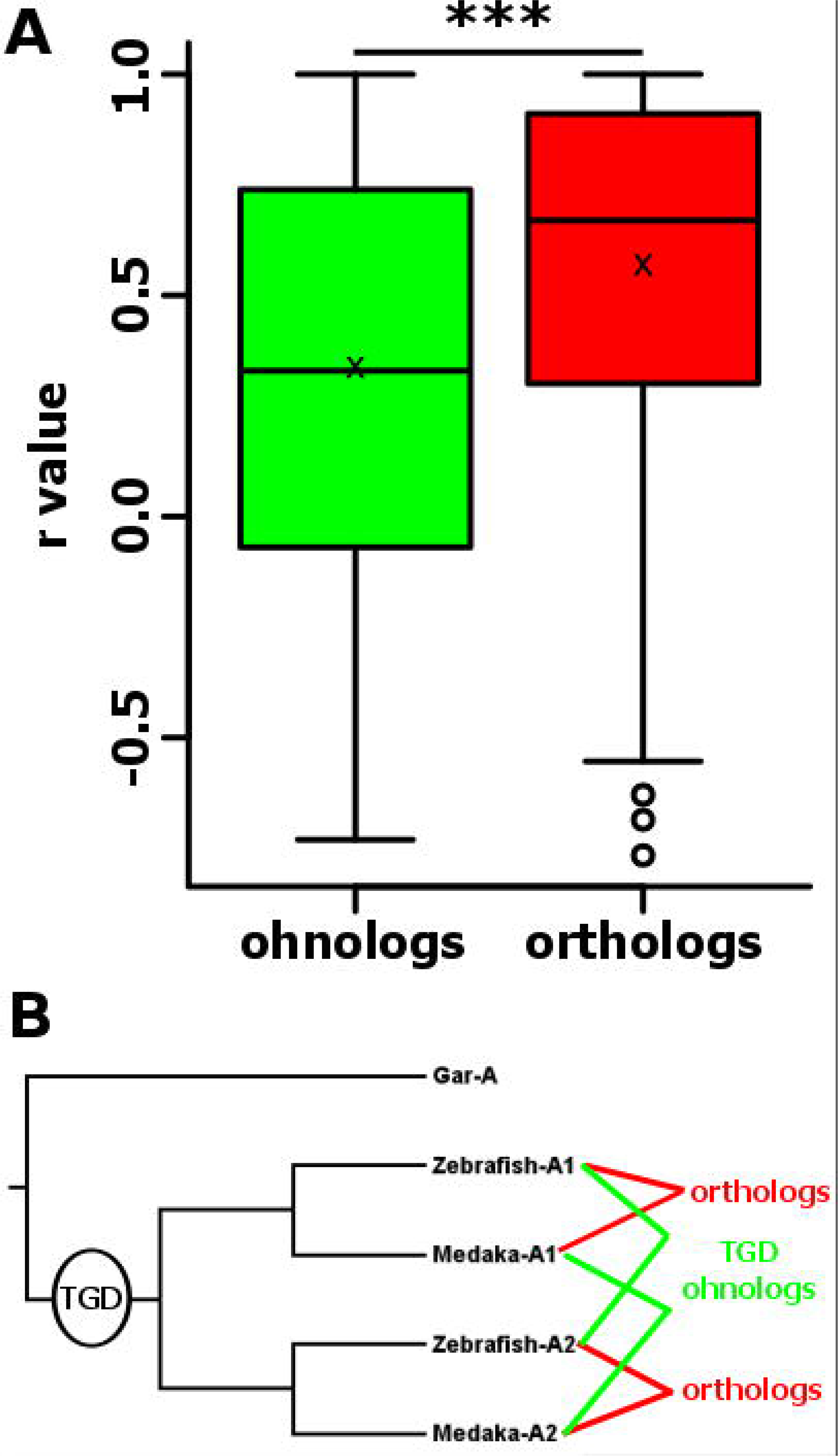
Distribution of expression pattern correlations between zebrafish and medaka TGD ohnologs and orthologs. For each boxplot (A), the black line, the black cross and the black circle represent the median, the mean and the outliers, respectively. (B) Relationships of ohnologs and orthologs.

#### Classification of TGD ohnologs based on correlation of tissue-specific expression and level of expression

The 774 pairs of TGD ohnologs shared between zebrafish and medaka (Fig. 1A, orange/purple intersection) were classified based on the correlation of expression patterns of ohnolog-1 and ohnolog-2 and differences in levels of expression between the two genes of each ohnolog pair (Fig. 6). Among the 774 pairs of genes analyzed in zebrafish and medaka, 44.8% belonged to the same category in both species, including 8.2% in the HCSE (High Correlation, Similar Expression) category, 3.0% in the HCSE (High Correlation, Differential Expression) category, 19.3% in the HCSE (No Correlation, Similar Expression) category, and 14.4% in the HCDE (No Correlation, Differential expression) category.

**Figure 6:**
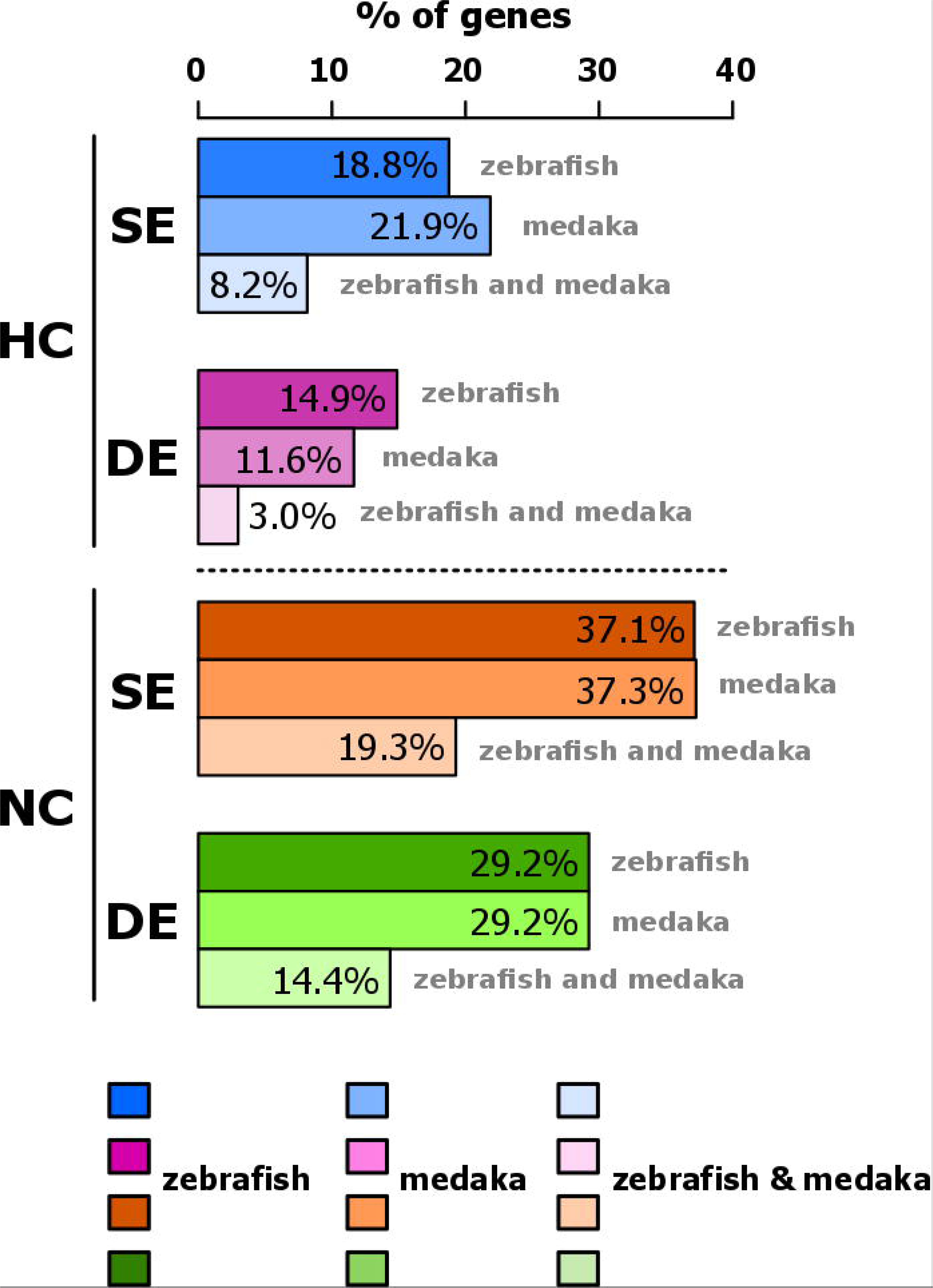
Expression of conserved TGD ohnologs in zebrafish and medaka reveals four classes of genes based on the conservation of their expression profiles. Determination of four groups of ohnologs based on (i) correlation between their expression patterns (HC: high correlation, p > 0.05; NC: no correlation, p < 0.05, Pearson’s correlation test), and (ii) their relative expression levels (SE: similar expression levels, p > 0.05; DE: different expression levels, p < 0.05, Student’s t-test). The percentage of genes present in each class (HCSE, HCDE, NCSE, and NCDE) is shown for zebrafish, medaka, and for genes belonging in the same class for the two species.

A large majority of genes were not significantly correlated (NC) with a total of 66.3% (37.1+29.2) and 66.5% (37.3+29.2) in the NC category for zebrafish and medaka, respectively. This high proportion of non-correlated genes was also observed for genes present in the same category in both zebrafish and medaka, with 33.7% (19.3+14.4) NC genes and 11.2% (8.2+3.0) of highly correlated (HC) genes (Figure 6).

In the NC category, a significantly higher proportion of genes exhibiting a similar level of expression (NCSE) was observed in both species (Fig. 6). In zebrafish, 37.1% and 29.2% of analyzed genes were classified into NCSE and NCDE categories, respectively. In medaka, 37.3% and 29.2% of analyzed genes were classified into NCSE and NCDE categories, respectively (Fig. 6). A similar trend was observed for the genes belonging to the same category in both zebrafish and medaka (Fig. 6).

For highly correlated (HC) genes, a higher proportion of genes exhibiting a similar level of expression (HCSE) was observed in both species, although this difference was not significant in zebrafish. In zebrafish, 18.8% and 14.9% of analyzed genes were classified into HCSE and HCDE categories, respectively (Fig. 6). In medaka, 21.9 % and 11.6% of analyzed genes were classified into HCSE and HCDE categories, respectively (Fig. 6). A higher proportion of genes exhibiting a similar level of expression (HCSE), in comparison to NCDE genes) was also observed for the genes belonging to the same category in both zebrafish and medaka (Fig. 6).

#### Evolution of TGD ohnolog expression in zebrafish and medaka

When looking at expression patterns of duplicated genes in different zebrafish and medaka tissues, HCSE genes were expressed predominantly in brain, bones, testis, and embryos (Fig. 7). This pattern was also observed for genes belonging to the HCSE group in both species (Fig. 7, central panel). Genes found in the HCSE category included zebrafish *bicd1* (Fig. 7, see also Fig. 2).

**Figure 7:**
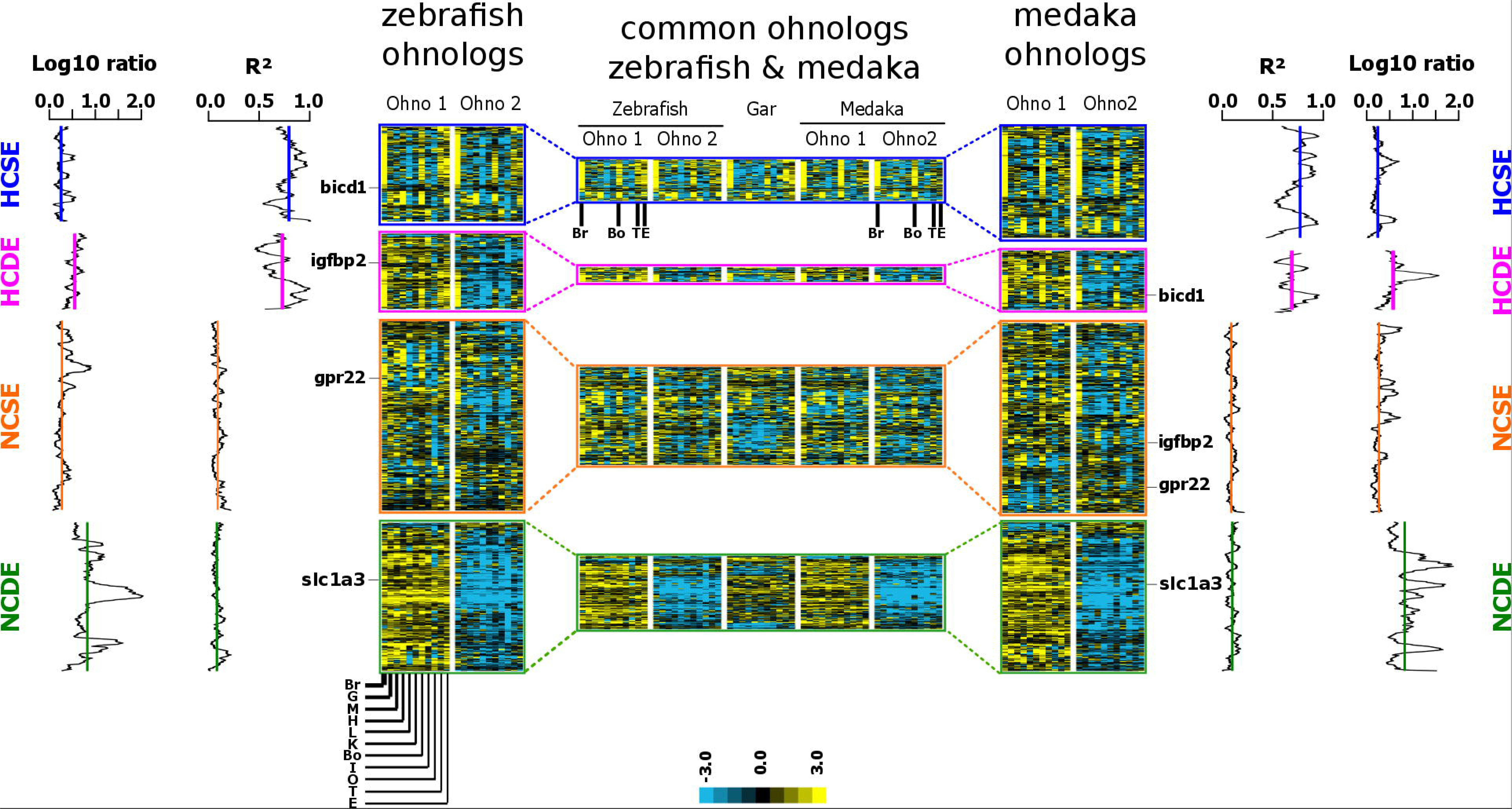
Expression of conserved TGD ohnologs in zebrafish and medaka. Heat maps of expression profiles of zebrafish (left panel) and medaka (right panel) ohnologs across 11 tissues (brain, gills, heart, muscle, liver, kidney, bone, intestine, ovary, testis and embryo). Correlations (R^2^) and log10 ratio of average expression between the two ohnologs are shown for genes belonging to HCSE, HCDE, NCSE, and NCDE categories as defined in Figure 6. The central panel shows the expression profiles of gar genes for which the two corresponding ohnologs are present in both zebrafish and medaka. All expression profiles were generated by hierarchical clustering analysis. Expression levels of both ohnologs were normalized and median-centered to highlight differences in relative levels of expression among both ohnologous genes in zebrafish and medaka separately and also (central panel) among common ohnolog pairs in zebrafish and medaka.

For ohnolog pairs that did not show significant tissue expression pattern correlations between the two paralogs (NCSE and NCDE categories, Fig 6), clustering analysis did not reveal any specific tissues exhibiting a predominant expression. For TGD ohnologs in which one of the two genes was significantly underexpressed in comparison to its ohnolog in both zebrafish and medaka (HCDE and NCDE groups, Fig. 7 central panel), a similar pattern was observed for orthologs (i.e. when zebrafish ohnolog-1 is overexpressed, its ortholog - medaka ohnolog-1- is overexpressed relative to ohnolog-2).

## Discussion

In addition to two whole-genome duplication events that occurred at the root of the vertebrate lineage (VGD1 and VGD2, Dehal et al. 2005; Nakatani et al. 2007; Canestro et al. 2009), teleosts experienced a third round of whole genome duplication (TGD, teleost-specific genome duplication)(Amores 1998; Postlethwait 1999; taylor 2003; jaillon 2004). The TGD occurred after the divergence between the holostean and teleostean lineages about 320-350 million years ago depending on the estimate (Hoegg et al., 2004; Amores et al., 2011). The recent publication of the gar genome sequence supports evidence that holosteans form a monophyletic lineage gathering gars and bowfin (Near et al., 2012; Betancur-R et al., 2013; Broughton et al., 2013; Faircloth et al., 2013; Braasch et al., 2016). The concomitant release of tissue-specific gene repertoires of holostean and teleostean species originating from our PhyloFish database project (Pasquier et al., 2016) offered the opportunity to cast new light on the fate of gene expression after TGD in teleost using spotted gar, medaka and zebrafish. These two model teleost species were selected because of their relatively distant position in the teleost tree of life (divergence estimate is 275 Mya (Near et al., 2012) and occurred relatively early within the teleosts supercohort Clupeocephalan), while gar was chosen as an ‘unduplicated’ holostean, pre-TGD-diverging reference. It should also be pointed out that genomic data from all three species was necessary to unambiguously infer orthology and paralogy relationships using conserved synteny information. The use of species with a well-assembled genome, such as gar, zebrafish, and medaka, was therefore required to generate the largest possible, yet stringent gene dataset for our analysis. In our study, we were able to use as much as 51.5% of annotated zebrafish genes and 60.4% of annotated medaka genes for which we could unambiguously identify a ohnolog-pair or singleton status with regard to the TGD and for which we could identified a single gar ortholog. According to Ensembl74, 18,328 protein-coding genes were annotated in the spotted gar genome (Braasch et al., 2016). In the present study, we used about 75% (13,734) of these gar genes and corresponding orthologs and co-orthologs in medaka and zebrafish. To our knowledge, the present work and the recent publication of the spotted gar genome (Braasch et al., 2016) correspond to one of the first genome-wide analysis of gene expression after a vertebrate WGD using such a high percentage of known genes.

After WGD by autopolyploidy, the daughter genes of each ancestral gene would be identical in terms of both coding and regulatory regions, and thus their functions would be fully redundant. In the case of allopolyploidy, however, the corresponding duplicates would be highly similar but in some cases might not be identical. The assumption of identical or highly similar functions implies that selective pressure should be lowered on both genes of each ohnologous pair. In principle, WGD gene pairs could undergo one or more of several different fates: (i) one of the duplicates could be lost (nonfunctionalization); (ii) both duplicates could be retained almost unchanged in expression; (iii) both duplicates could evolve different in ways that result in the partitioning of the ancestral function - qualitatively or quantitatively - between the two duplicates (subfunctionalization); and finally (iv) one of the duplicate genes could acquire a new function (neofunctionalization). Those categories are simplified and multiple processes could affect the evolution of the same duplicated gene pair simultaneously or successively.

### Loss of one of the duplicate genes is the most common fate after the TGD

In our study, a large majority (87% (10,415 out 12,021 in zebrafish, see Fig.1) and 88% (9,265 out of 10,580 in medaka, see Fig. 1)) of genes were retained only as singletons after the TGD, while less than 15% (12-13%, see Fig. 1) were kept in duplicate. These observations are consistent with previous studies that reported 5% and 20 % of genes retained in duplicates in pufferfish (Jaillon et al., 2004) and zebrafish (Postlethwait et al., 2000, 2004), respectively. Our study was performed on a larger and perhaps less selected gene data set, thus leading to a more precise estimation of the frequency of ohnologs in teleosts. Together these observations confirm that non-functionalization (i.e. loss of one of the duplicated copy) is the most common fate of duplicate genes after TGD.

When considering genes present in singletons in both species, we observed that 70% of zebrafish singletons were also singletons in medaka and 79% of medaka singletons were singletons in zebrafish. The large proportion (>70%) of singletons shared by medaka and zebrafish - two lineages that diverged more than 250 Mya (Near et al., 2012) - indicates that the loss of one of the two duplicated copies occurred shortly after TGD (i.e. in the 70 to 100 My after the TGD). We found that at least 15.6% (2,147/13,734) of genes were still present as TGD ohnologs in the last common clupeocephalan ancestor of zebrafish and medaka. These estimates are in agreement with the study of Inoue and coworkers which estimated that 70 to 80 % of the gene duplicates were lost during the 60 My following the TGD (Inoue et al., 2015). Together, these results are fully consistent with studies in Eukaryotes predicting that the vast majority of duplicated genes are silenced within a few million years after duplication (Lynch and Conery, 2000). Importantly, our derived lists here are likely an underestimate of the actual number of retained TGD ohnologs in general due to our stringent filtering by gene phylogeny topology and conserved synteny as well as the use of only two teleost species.

### The average expression of ohnolog pairs resembles the ancestral pattern

The present study shows that the average expression of pairs of ohnologs is more correlated to the expression pattern of their gar ortholog than are singletons or than is each ohnolog taken separately. The higher correlation observed for pairs of ohnologs, in comparison to each ohnolog taken separately, is not surprising because each member of a pair of ohnologs is likely to behave differently when undergoing neofunctionalization and/or subfunctionalization. The higher correlation of pairs of ohnologs to gar genes compared to the correlation of singletons to their gar ortholog is, in contrast, more surprising. This finding could be due, at least in part, to the marked difference in the number of genes in the different categories (close to 10,000 for singletons and below 2,000 for ohnologs). These observations are more likely to mean, however, that pairs of ohnologs when averaged are more likely than singletons to reflect the ancestral expression patterns, which can be inferred from expression patterns in the slow evolving spotted gar. This result would be observed if ancestral functions partitioned between duplicates (subfunctionalization), if subfunctionalization had already occurred before nonfunctionalization, and if singletons are likely to have evolved different functions in different teleost lineages after lineages diverged.

### Evolution of gene expression when both duplicates are retained

After TGD, genes are retained in duplicates in about 12-13% of our gene set. When both ohnologs are retained by one species, both members of the pairs also tend to be found in the other species, zebrafish (48%) or medaka (59%). This result suggests that fixation of both copies occurred relatively rapidly after the TGD, consistent with existing literature (Force et al., 1999; Lynch and Conery, 2000), even though the subsequent loss of one copy happened in a lineage-dependent manner during and/or after the speciation event leading to the zebrafish and medaka lineages.

### Expression patterns of ohnologs are poorly correlated

Here we show that, following the TGD, the expression pattern of one member of a pair of ohnologs is poorly correlated with the expression pattern of the other ohnolog. In contrast, a much higher correlation is observed between the expression patterns of orthologs. This observation suggests that orthologs between zebrafish and medaka tend to retain similar functions more often than ohnologs within a species. This finding indicates that, when retained in duplicates, the majority of gene pairs undergo a significant divergence in the expression patterns of the two members of the pair, most likely associated with an evolution of their functions.

#### Genes undergoing progressive silencing

In both zebrafish and medaka, we observed that a small proportion (less than 15%, Figure 6) of dual conserved ohnologs exhibit highly correlated expression (i.e. ohnolog-1 highly correlated to ohnolog-2, or HC) with a lower expression of one of the two copies (DE). This result is in striking contrast with a study in rainbow trout (Order Salmoniformes, family Salmonidae, a salmonid species in which a more recent WGD - the salmonid-specific genome duplication, SaGD-occurred 100 Mya) reporting over 30% of SaGD ohnologs falling in this HCDE category (Berthelot et al., 2014). This indicates that, among ohnologs, only a small fraction of genes are undergoing silencing in medaka and zebrafish. It is, however, unknown if this fraction (i.e. ohnologs undergoing silencing) is the remnant of progressive silencing of one duplicate following the TGD or if it corresponds to the ‘natural’ background gene silencing observed during and/or after speciation. The low frequency (3%, Figure 6) of genes of this category (HCDE) found in both zebrafish and medaka suggests that this gene silencing process has mostly occurred in a species dependent manner and would therefore favor the second hypothesis.

#### Genes undergoing neofunctionalization or subfunctionalization

Our results clearly indicate that, after the TGD, a large majority of ohnologs undergo major changes in expression pattern and/or levels. While subfunctionalization and neofunctionalization processes may occur simultaneously or consecutively, we were able to identify clear-cut cases of subfunctionalization and neofunctionalization that were previously unknown (listed in Supplemental file 3). Among the remarkable cases of neofunctionalization is *solute carrier family 1 member 3* (*slc1a3*), which exhibited a novel expression in the liver in both zebrafish and medaka, while *bicaudal D homolog 1* (*bicd1*) exhibited a novel expression in the medaka heart. Similarly, *G protein-coupled receptor 22* (*gpr22*) exhibited clear subfunctionalization of TGD ohnologs in both teleosts, as did medaka *insulin-like growth factor binding protein 2* (*igfbp2*). Clear cases of subfunctionalization and neofunctionalization included cases of gene expression changes that occurred before or after the divergence of zebrafish and medaka lineages. The number of neofunctionalization and subfunctionalization examples reported here are, however, too low to conclude whether or not most neofunctionalization and subfunctionalization events occurred soon after the TGD or later (i.e. during/after divergence of zebrafish and medaka lineages). The percentage of non-correlated profiles with similar expression levels (NCSE category in Figures 6 and 7) common to both species, or in contrast present in a single species, suggests that changes in gene expression including neofunctionalization and subfunctionalization events occurred not only rapidly after the TGD but also later in teleost evolution. This conclusion would be consistent with existing evidence that asymmetric neofunctionalization or subfunctionalization of TGD gene pairs could drive long term diversification and speciation events in teleosts.

In addition, we note that after TGD, genomic rearrangements can relocate a TGD paralog from of its TGD paralogon into a new, non-syntenic genomic environment. Conceivably, such relocated TGD paralogs might be particularly prone to expressional neofunctionalization due to novel gene regulatory inputs in their genomic vicinity. Such cases of relocated TGD paralogs, however, are indistinguishable from other types of gene duplication given our stringent filtering based on conserved synteny support for TGD ohnology, which may have underestimated the occurrence of neofunctionalization among TGD paralog pairs. Finally, a recent work by Lien and coworkers in Atlantic salmon reported far more instances of neofunctionalization than subfunctionalization (Lien et al., 2016). This conclusion is consistent with the number of clear neofunctionalization and subfunctionalization cases reported in the present study. In contrast, our data also suggest that the average expression of duplicated teleost genes often approximate the patterns and levels of expression for gar genes, consistent with subfunctionalization (Braasch et al., 2016). Together, these results indicated that quantifying the respective occurrence of neofunctionalization and subfunctionalization following WGD is complex and will requires further investigation and novel tools and approaches.

### Twenty percent of duplicates are retained almost unchanged possibly due to gene dosage effects on gene expression

In both zebrafish and medaka, approximately 20% (18.8-21.9, Figure 6) of ohnologs exhibit a highly correlated expression profile and similar expression levels (HCSE). In rainbow trout, a similar frequency was observed for SaGD ohnologs (Berthelot et al., 2014). Together, these observations indicate that in approximately 20% of the cases, retaining a similar expression of duplicated genes is apparently not strongly selected against. As previously discussed by Force and coworkers (Force et al., 1999), gene dosage requirements participate in the evolution of gene expression following duplication. In the present study, we observed that ohnologs with correlated profiles and similar expression levels are especially abundant in brain, bones, embryo, and testis. This finding would suggest that nervous system-related functions, mineralization, and male reproduction are processes in which duplicates are retained relatively unchanged, suggesting that expression dosage may be particularly important for those organs.

## Conclusion

Together, our data show that most of the TGD duplicates acquired their current status (loss of one duplicate gene or retention of both ohnologs) shortly after the TGD and before the divergence of zebrafish and medaka lineages, which separated about 275 Mya. Results demonstrate that in both zebrafish and medaka, the loss of one of the duplicate genes is the most common fate after TGD with a probability of about 80%. In addition, results provided evidence that the fate of duplicate genes after TGD, including subfunctionalization, neofunctionalization, or retention of two ‘almost similar’ copies occurred not only rapidly after the TGD but continued later during evolution consistent with potential roles in long-term diversification and presumably adaptive radiation. Finally, analysis revealed novel cases of subfunctionalization and neofunctionalization that further illustrate the importance of these two processes on long-term retention of duplicated genes after TGD.

## Acknowledgements

The authors thank the BROAD Institute for generating the spotted gar genome assembly and MGX-Montpellier GenomiX for performing RNA-seq, and Allyse Ferrara and Quenton Fontenot (Nicholls State University) for sampling spotted gar tissues.

## Conflicts of interest

The authors have nothing to declare.

## Supplemental Material

**Supplemental File 1: Criteria used for detection of conserved neo- and subfunctionalized TGD ohnologs**. Pearson correlation coefficient (r) was used as criteria to identify neo- and subfunctionalized TGD ohnologs in both zebrafish and medaka.

**Supplemental File 2: Correspondance tables between gar genes and their orthologs in zebrafish and medaka**. Each genes are listed as their Ensembl unique identifiers (http://www.ensembl.org/index.html). Gar genes are partitioned according the diagramme of Figure 1.

**Supplemental File 3: Table of conserved neofunctionalized and subfunctionalized ohnologs in zebrafish and medaka, based on expression pattern comparisons with their gar orthologs**. Neo- and subfunctionalized TGD ohnologs are partitioned into two groups, i.e. early and late, depending on the occurrence of the neo- or subfunctionalization processes in both or in only one of the two teleost species, respectively.

## Notes

This work was supported by ANR under grant agreement #ANR-10-GENM-017 (PhyloFish) to JB; NIH grants R01 OD011116 (alias R01 RR020833) and R24 OD01119004 (J.H.P.); a Feodor Lynen Fellowship from the Alexander von Humboldt Foundation and the Volkswagen Foundation Initiative Evolutionary Biology, grant I/84 815 (I.B.).

## References

Amores A, Catchen J, Ferrara A, Fontenot Q, Postlethwait JH. 2011. Genome evolution and meiotic maps by massively parallel DNA sequencing: spotted gar, an outgroup for the teleost genome duplication. Genetics [Internet] 188:799–808. Available from: http://www.pubmedcentral.nih.gov/articlerender.fcgi?artid=3176089&tool=pmcentrez&rendertype=abstract

Amores A, Force A, Yan YL, Joly L, Amemiya C, Fritz A, Ho RK, Langeland J, Prince V, Wang YL, Westerfield M, Ekker M, Postlethwait JH. 1998. Zebrafish hox clusters and vertebrate genome evolution. Science (80- ) 282:1711–1714.

Anders S, Huber W. 2010. Differential expression analysis for sequence count data. Genome Biol [Internet] 11:R106. Available from: http://www.ncbi.nlm.nih.gov/pubmed/20979621

Berthelot C, Brunet F, Chalopin D, Juanchich A, Bernard M, Noël B, Bento P, Da Silva C, Labadie K, Alberti A, Aury J-M, Louis A, Dehais P, Bardou P, Montfort J, Klopp C, Cabau C, Gaspin C, Thorgaard GH, Boussaha M, Quillet E, Guyomard R, Galiana D, Bobe J, Volff J-N, Genêt C, Wincker P, Jaillon O, Roest Crollius H, Guiguen Y. 2014. The rainbow trout genome provides novel insights into evolution after whole-genome duplication in vertebrates. Nat Commun [Internet] 5:3657. Available from: http://www.nature.com/ncomms/2014/140422/ncomms4657/full/ncomms4657.html

Betancur-R R, Broughton RE, Wiley EO, Carpenter K, López JA, Li C, Holcroft NI, Arcila D, Sanciangco M, Cureton Ii JC, Zhang F, Buser T, Campbell MA, Ballesteros JA, Roa-Varon A, Willis S, Borden WC, Rowley T, Reneau PC, Hough DJ, Lu G, Grande T, Arratia G, Ortí G. 2013. The tree of life and a new classification of bony fishes. PLoS Curr [Internet] 5. Available from: http://www.pubmedcentral.nih.gov/articlerender.fcgi?artid=3644299&tool=pmcentrez&rendertype=abstract

Braasch I, Gehrke AR, Smith JJ, Kawasaki K, Manousaki T, Pasquier J, Amores A, Desvignes T, Batzel P, Catchen J, Berlin AM, Campbell MS, Barrell D, Martin KJ, Mulley JF, Ravi V, Lee AP, Nakamura T, Chalopin D, Fan S, Wcisel D, Cañestro C, Sydes J, Beaudry FEG, Sun Y, Hertel J, Beam MJ, Fasold M, Ishiyama M, Johnson J, Kehr S, Lara M, Letaw JH, Litman GW, Litman RT, Mikami M, Ota T, Saha NR, Williams L, Stadler PF, Wang H, Taylor JS, Fontenot Q, Ferrara A, Searle SMJ, Aken B, Yandell M, Schneider I, Yoder JA, Volff J-N, Meyer A, Amemiya CT, Venkatesh B, Holland PWH, Guiguen Y, Bobe J, Shubin NH, Di Palma F, Alföldi J, Lindblad-Toh K, Postlethwait JH. 2016. The spotted gar genome illuminates vertebrate evolution and facilitates human-teleost comparisons. Nat Genet [Internet]. Available from: http://www.ncbi.nlm.nih.gov/pubmed/26950095

Broughton RE, Betancur-R R, Li C, Arratia G, Ortí G. 2013. Multi-locus phylogenetic analysis reveals the pattern and tempo of bony fish evolution. PLoS Curr [Internet] 5. Available from: http://www.pubmedcentral.nih.gov/articlerender.fcgi?artid=3682800&tool=pmcentrez&rendertype=abstract

Canestro C, Catchen JM, Rodriguez-Mari A, Yokoi H, Postlethwait JH. 2009. Consequences of lineage-specific gene loss on functional evolution of surviving paralogs: ALDH1A and retinoic acid signaling in vertebrate genomes. PLoSGenet 5:e1000496-.

Catchen JM, Conery JS, Postlethwait JH. 2009. Automated identification of conserved synteny after whole-genome duplication. Genome Res 19:1497–1505.

Clarke JT, Lloyd GT, Friedman M. 2016. Little evidence for enhanced phenotypic evolution in early teleosts relative to their living fossil sister group. Proc Natl Acad Sci [Internet] 113:11531–11536. Available from: http://www.ncbi.nlm.nih.gov/pubmed/27671652

Dehal P, Boore JL. 2005. Two rounds of whole genome duplication in the ancestral vertebrate. PLoSBiol 3:e314-.

Eisen MB, Spellman PT, Brown PO, Botstein D. 1998. Cluster analysis and display of genome-wide expression patterns. Proc Natl Acad Sci U S A 95:14863–14868.

Faircloth BC, Sorenson L, Santini F, Alfaro ME. 2013. A Phylogenomic Perspective on the Radiation of Ray-Finned Fishes Based upon Targeted Sequencing of Ultraconserved Elements (UCEs). PLoS One [Internet] 8:e65923. Available from: http://dx.plos.org/10.1371/journal.pone.0065923

Force A, Lynch M, Pickett FB, Amores A, Yan YL, Postlethwait J. 1999. Preservation of duplicate genes by complementary, degenerative mutations. Genetics 151:1531–1545.

Glasauer SMK, Neuhauss SCF. 2014. Whole-genome duplication in teleost fishes and its evolutionary consequences. Mol Genet Genomics [Internet] 289:1045–60. Available from: http://www.ncbi.nlm.nih.gov/pubmed/25092473

He X, Zhang J. 2005. Rapid subfunctionalization accompanied by prolonged and substantial neofunctionalization in duplicate gene evolution. Genetics [Internet] 169:1157–64. Available from: http://www.ncbi.nlm.nih.gov/pubmed/15654095

Hoegg S, Brinkmann H, Taylor JS, Meyer A. 2004. Phylogenetic timing of the fish-specific genome duplication correlates with the diversification of teleost fish. J MolEvol 59:190–203.

Inoue J, Sato Y, Sinclair R, Tsukamoto K, Nishida M. 2015. Rapid genome reshaping by multiple-gene loss after whole-genome duplication in teleost fish suggested by mathematical modeling. Proc Natl Acad Sci U S A [Internet] 112:14918–23. Available from: http://www.ncbi.nlm.nih.gov/pubmed/26578810

Jaillon O, Aury J-M, Brunet F, Petit J-L, Stange-Thomann N, Mauceli E, Bouneau L, Fischer C, Ozouf-Costaz C, Bernot A, Nicaud S, Jaffe D, Fisher S, Lutfalla G, Dossat C, Segurens B, Dasilva C, Salanoubat M, Levy M, Boudet N, Castellano S, Anthouard V, Jubin C, Castelli V, Katinka M, Vacherie B, Biémont C, Skalli Z, Cattolico L, Poulain J, De Berardinis V, Cruaud C, Duprat S, Brottier P, Coutanceau J-P, Gouzy J, Parra G, Lardier G, Chapple C, McKernan KJ, McEwan P, Bosak S, Kellis M, Volff J-N, Guigó R, Zody MC, Mesirov J, Lindblad-Toh K, Birren B, Nusbaum C, Kahn D, Robinson-Rechavi M, Laudet V, Schachter V, Quétier F, Saurin W, Scarpelli C, Wincker P, Lander ES, Weissenbach J, Roest Crollius H. 2004. Genome duplication in the teleost fish Tetraodon nigroviridis reveals the early vertebrate proto-karyotype. Nature [Internet] 431:946–57. Available from: http://www.ncbi.nlm.nih.gov/pubmed/15496914

Kinsella RJ, Kähäri A, Haider S, Zamora J, Proctor G, Spudich G, Almeida-King J, Staines D, Derwent P, Kerhornou A, Kersey P, Flicek P. 2011. Ensembl BioMarts: a hub for data retrieval across taxonomic space. Database (Oxford) [Internet] 2011:bar030. Available from: http://www.pubmedcentral.nih.gov/articlerender.fcgi?artid=3170168&tool=pmcentrez&rendertype=abstract

Li H, Durbin R. 2009. Fast and accurate short read alignment with Burrows-Wheeler transform. Bioinformatics 25:1754–1760.

Li H, Handsaker B, Wysoker A, Fennell T, Ruan J, Homer N, Marth G, Abecasis G, Durbin R, 1000 Genome Project Data Processing Subgroup. 2009. The Sequence Alignment/Map format and SAMtools. Bioinformatics [Internet] 25:2078–2079. Available from: http://www.ncbi.nlm.nih.gov/pubmed/19505943

Lien S, Koop BF, Sandve SR, Miller JR, Matthew P, Leong JS, Minkley DR, Zimin A, Grammes F, Grove H, Gjuvsland A, Walenz B, Hermansen RA, Schalburg K Von, Rondeau EB, Genova A Di, Samy JKA, Vik JO. 2016. The Atlantic salmon genome provides insights into rediploidization. Nature 533.

Lynch M, Conery JS. 2000. The evolutionary fate and consequences of duplicate genes. Science (80- ) 290:1151–1155.

Nakatani Y, Takeda H, Kohara Y, Morishita S. 2007. Reconstruction of the vertebrate ancestral genome reveals dynamic genome reorganization in early vertebrates. Genome Res [Internet] 17:1254–1265. Available from: http://www.genome.org/cgi/doi/10.1101/gr.6316407

Near TJ, Eytan RI, Dornburg A, Kuhn KL, Moore JA, Davis MP, Wainwright PC, Friedman M, Smith WL. 2012. Resolution of ray-finned fish phylogeny and timing of diversification. Proc Natl Acad Sci U S A [Internet] 109:13698–703. Available from: http://www.pubmedcentral.nih.gov/articlerender.fcgi?artid=3427055&tool=pmcentrez&rendertype=abstract

Ohno S. 1970. Evolution by Gene Duplication. Berlin, Heidelberg: Springer Berlin Heidelberg. Available from: http://link.springer.com/10.1007/978-3-642-86659-3

Pasquier J, Cabau C, Nguyen T, Jouanno E, Severac D, Braasch I, Journot L, Pontarotti P, Klopp C, Postlethwait JH, Guiguen Y, Bobe J. 2016. Gene evolution and gene expression after whole genome duplication in fish: the PhyloFish database. BMC Genomics [Internet] 17:368. Available from: http://www.ncbi.nlm.nih.gov/pubmed/27189481

Postlethwait J, Amores A, Force A, Yan YL. 1999. The zebrafish genome. Methods Cell Biol [Internet] 60:149–63. Available from: http://www.ncbi.nlm.nih.gov/pubmed/9891335

Postlethwait J, Ruotti V, Carvan MJ, Tonellato PJ. 2004. Automated analysis of conserved syntenies for the zebrafish genome. Methods Cell Biol [Internet] 77:255–71. Available from: http://www.ncbi.nlm.nih.gov/pubmed/15602916

Postlethwait JH, Woods IG, Ngo-Hazelett P, Yan YL, Kelly PD, Chu F, Huang H, Hill-Force A, Talbot WS. 2000. Zebrafish comparative genomics and the origins of vertebrate chromosomes. Genome Res [Internet] 10:1890–902. Available from: http://www.ncbi.nlm.nih.gov/pubmed/11116085

Santini F, Harmon LJ, Carnevale G, Alfaro ME. 2009. Did genome duplication drive the origin of teleosts? A comparative study of diversification in ray-finned fishes. BMC Evol Biol [Internet] 9:194. Available from: http://bmcevolbiol.biomedcentral.com/articles/10.1186/1471-2148-9-194

Taylor JS, Braasch I, Frickey T, Meyer A, Van de PY. 2003. Genome duplication, a trait shared by 22000 species of ray-finned fish. Genome Res 13:382–390.

